# Charting the Silent Signals of Social Gaze: Automating Eye Contact Assessment in Face-to-Face Conversations

**DOI:** 10.1101/2024.08.28.610064

**Authors:** Ralf Schmälzle, Nolan T. Jahn, Gary Bente

## Abstract

Social gaze is a crucial yet often overlooked aspect of nonverbal communication. During conversations, it typically operates subconsciously, following automatic co-regulation patterns. However, deviations from typical patterns, such as avoiding eye contact or excessive gazing, can significantly affect social interactions and perceived relationship quality. The principles and effects of social gaze have intrigued researchers across various fields, including communication science, social psychology, animal biology, and psychiatry. Despite its significance, research in social gaze has been limited by methodological challenges in assessing eye movements and gaze direction during natural social interactions. To address these obstacles, we have developed a new approach combining mobile eye tracking technology with automated analysis tools. In this paper, we introduce, validate, and apply a pipeline for recording and analyzing gaze behavior in dyadic conversations. We present a sample study where dyads engaged in two types of interactions: a get-to-know conversation and a conflictual conversation. Our new analysis pipeline corroborated previous findings, such as people directing more eye gaze while listening than talking, and gaze typically lasting about three seconds before averting. These results demonstrate the potential of our methodology to advance the study of social gaze in natural interactions.

## Introduction

Imagine waking up in a world where eye contact has become impossible: Conversations have become hollow, devoid of the unspoken understanding between the eyes of speakers and listeners. Interactions with partners, family, and friends lack the closeness and intimacy that comes with eye contact. In business meetings, colleagues struggle to gauge sincerity, leading to an erosion of trust. As this scenario illustrates, we may often take social gaze for granted, but this capacity to send and receive signals with the eyes is central to communication (Patterson, 1982; Kleinke, 1986). If eye contact became impossible, we may find ourselves adrift in a sea of words, but miss the coordinating and connecting functions provided by social gaze. Indeed, this is actually a real risk in zoom meetings and increasingly avatar-mediated communication, where technology fails to accurately represent gaze signals.

Given its importance for communication, researchers have long strived to examine the fine mechanisms and strong effects of social gaze and associated nonverbal behaviors (Duncan and Fiske, 1977; Mehrabian, 1971). However, these efforts have been plagued by operational challenges, particularly the labor-intensive and potentially imprecise nature of video-based interaction recordings and manual analysis methods. Thus, there is a need to improve methodologies to unlock the next stage of the oretical research on nonverbal communication. This is the goal of the current work.

Overview: In Part I of this manuscript, we discuss the role of social gaze for communication and locate its relevance within the broader literature on nonverbal communication. In Part II, we introduce and validate an automated pipeline for measuring social gaze. Then, in Part III, we apply the pipeline to an example dataset of naturalistic conversations under two social conditions, a get-to-know and a managerial interaction. We provide quantitative metrics that characterize the eye-contact and compare the influence of conversational (speaker vs listener) and social (manager vs. employee) roles. Finally, we discuss results and broader implications.

## Background

### Role of Social Gaze in Communication

The eyes have long been recognized as a vital component of social interaction (Kendon, 1967; Duncan and Fiske, 1977; Pfeiffer et al., 2013). Social gaze behavior serves multiple functions related to signaling attention, coordinating the conversation, and regulating interpersonal distance (Patterson, 1982). Social gaze encompasses directed and averted gaze (i.e. looking at or away from the partner) as well as pivotal moments of eye-contact (Kleinke, 1986). In dyadic conversations, the partners’ eyes engage in an intricate dance that can convey a wealth of information, from intimacy to discomfort, confidence to submission, as well as cultural characteristics (Argyle and Cook,X 1976; Foddy, 1978; Cook, 1977).

Although studies of social gaze have been part and parcel of nonverbal communication research for decades, many open questions remain. In fact, even nonverbal textbooks rarely devote more than one to two pages to gaze and basic statistics of social gaze behavior are not well known (Burgoon, 1996; Mehrabian, 1971). In fact, it seems safe to say that most people will know more about the energy-efficiency of their cars or heating systems than quantitative descriptives of human social gaze. Two such facts about social gaze are that people tend to look more at the partner while listening and that the duration of directed gaze (looking at the eye/face of the interaction partner) is about 2-3 seconds (too short durations come across as erratic and too long durations will be interpreted as uncomfortable staring).

Recent years have seen an increase in research on social gaze. Work on social gaze and nonverbal communication more broadly was very popular in the 1970s-80s (Argyle and Cook, 1976; Burgoon and Buller, 1994; Cappella and Green, 1984; Duncan and Fiske, 1977), but faded in the 1990s and 2000s, likely as a result of the operational difficulties associated with collecting and analyzing data and the comparative ease with which data could be collected via survey and online research methods. Over the past years, however, there appears to be a resurgence of interest (Jongerius et al., 2020). Related work in cognitive psychology and visual perception has started to take an interest in examining dyadic gaze, above and beyond the long-standing occupation with face perception in more static settings (Macdonald and Tatler, 2013; Carlin and Calder, 2013; Risko et al., 2016). Coming more from a clinical orientation, a growing body of research aims to study gaze for diagnostic purposes (e.g. in autism), or to gain insight into the biological roots of abnormal gaze patterns (Pfeiffer et al., 2013; Georgescu et al., 2014). Finally, several studies have examined social gaze in developmental, applied, or technological contexts, from early parent-infant interactions (Hoehl et al., 2014), doctor-patient communication (Jongerius et al., 2021), linguistic turn-taking (Wohltjen and Wheatley, 2021), or interactions with embodied conversational agents (Bente et al., 2007; Georgescu et al., 2013).

### The Measurement Dilemma: Struggle to Capture and Analyze Social Gaze in Conversations

When reading classical research on social gaze, it is clear that both measurement and analysis methods suffered from major limitations regarding objectivity, reliability, and validity. For example, many studies relied on trained observers who were seated behind participants looking to see if the conversational partner is engaged in eye or face gaze (Argyle and Cook, 1976; Kleinke, 1986). Other work used video-recording, but often only from single person, not the entire dyad; video-based recordings were then analyzed by coders to determine whether people looked at each other. However, without the use of eye-tracking, the validity of this approach is questionable. Perhaps unsurprisingly, interobserver reliabilities were wide ranging during these studies, averaging around *κ* = 0.66 (Argyle and Cook, 1976; Kleinke, 1986). Some factors that impact reliability are the distance between the participants, the angles of the observers relative to the participants, and simply the conversation duration as longer durations can lead to coder wearout (Kleinke, 1986). Finally, to keep the already effortful human coding manageable, coders would often just assess every five seconds if eye gaze had been present or not. This coarse temporal granularity clearly fails to capture intricacies of gaze patterns (Argyle and Cook, 1976; Burgoon and Le Poire, 1999; Kleinke, 1986).

Additionally, the field operated with ambiguous definitions and operationalizations (Kleinke, 1986; Jongerius et al., 2021). For instance, although a definitional distinction can be made between directed/averted gaze (directed at/averted from the person as a whole), face gaze (gazing at the face) and eye-gaze (gazing at the eye), in practice the methods used (observation and video recordings without eye-tracking) were unable to afford such precision. Moreover, terminology varied especially when it comes to the term eye-contact, which has sometimes been used to refer simply to gazing to the eyes (individually), but other times to refer the reciprocal nature of this behavior (i.e. mutual eye-contact; Jongerius et al. (2021). Here, we will use the term eye-contact to refer to reciprocal gazing towards the face area of each conversation partner. This operationalization i) honors the reciprocal nature of this key social phenomenon, ii) it matches the resolution that current measurements afford, iii) corresponds with typical face-viewing behaviors (Peterson and Eckstein, 2012), and iv) aligns with peoples’ subjective experiences of being looked at (Rogers et al., 2019).

### Precision Meets Practicality: Innovative Tools for Quantifying Social Gaze

The advent of eye tracking technology has brought gaze research to the forefront in many fields - from commercial marketing to health communication to media psychology (Carter and Luke, 2020; Duchowski, 2002; Schmälzle et al., 2023; Mackert et al., 2013). However the rigid and immobile experimental conditions that conventional screen-based eye-trackers required made them less suited for interpersonal communication studies. When assessing eye gaze in a natural conversation though, both the participant’s head is mobile and so is the head of the interaction partner (target). This creates immense analytical difficulties not present in the oldest studies that relied on two vantage points for human observers, or in the studies that relied on immobile participants and static targets. As a result, only a handful of studies has even attempted to examine social gaze in naturalistic and dyadic settings that are relevant for communication, and all were published only recently (Ho et al., 2015; Wohltjen and Wheatley, 2021; Macdonald and Tatler, 2018; Jongerius et al., 2021; Valtakari et al., 2021; Vehlen et al., 2021).

Recent advancements in eye tracking technology and analysis capabilities are overcoming these methodological limitations. Specifically, mobile eye trackers can capture eye gaze in freely behaving participants, and computer vision algorithms can dynamically detect target the participants’ field of view (e.g. the face of an interaction partner, which can move around based on the observer’s as well as the target’s movements). With this, it is now possible to capture social gaze “in the wild” and - provided that two mobile eye-trackers are employed - to do so in interpersonal settings like natural conversations. For instance, Ho and colleagues (Ho et al., 2015) used two mobile eye-tracking devices to record social gaze in a dyadic gaming setting, although the specific system they used was still relatively bulky and unnatural-looking. However, the latest generations of mobile eye-tracking glasses, such as the Pupil-Labs invisible, the Tobii Pro, and to some degree even the increasingly popular devices by Meta or Google are far more ecologically valid. These mobile eye-tracker look like normal glasses and do not interfere with neither the participants behavior nor the observer’s social perceptions. Thus, with these mobile eye-trackers, it is possible to record gaze in natural social interactions.

Importantly though, many of the recent studies using these mobile eye-trackers still rely on human observers to manually annotate the recorded gaze data (Ho et al., 2015; Vehlen et al., 2021; Wohltjen and Wheatley, 2021). This quickly becomes a monumental task, as typical trackers record video at rates of 30 frames per second. Thus, even if interactions are only 5 minutes long, this requires annotating 18000 frames (2 interactants, 300 sec * 30 fps) per dyad. As a result, some work has begun to study automated methods for analyzing mobile eye tracking data (Jongerius et al., 2021), reporting generally favorable results. However, the rapid evolution of both the eye-tracking devices as well as advanced computer-vision/AI algorithms warrants further study.

In sum, this all highlights the opportunity for the field of communication to embrace new mobile eye-trackers in combination with automated analysis routines. In the remainder of this paper, we introduce a new approach to study social gaze in dyadic conversations. We first introduce our measurement approach, which uses two mobile eye-trackers to capture participants’ gaze in naturally running social interactions. We then describe the pipeline for analyzing gaze (as well as accompanying speech) data recorded from social interactions. Demonstrating the feasibility of this approach requires answering two basic questions:

The first question is whether mobile eye-trackers precise enough to capture eye-contact (i.e. mutual/reciprocal gaze towards the face, see above)? The second question is whether automated analysis routines are able to accurately detect eye-contact? We present two studies that answer these questions, providing evidence that we can - and in fact should - use mobile eye-trackers to study social gaze.

Finally, we then apply the developed and validated pipeline to a dataset of 78 dyadic interactions (39 dyads, 78 participants, all performing two interactions) to characterize social gaze dynamics and demonstrate the potential of this approach.

### Validating a Computerized Pipeline for Individual and Dyadic Social Gaze Analysis

In this section, we describe the approach to measure eyegaze in naturalistic social interactions and provide evidence for its validity. In the accompanying online materials, we offer all code and a tutorial dataset allowing others to reproduce our results and apply the pipeline to their own data, including data collected with different eye-tracking hardware.

### Step-By-Step-Overview of Dyadic Interaction Data Recording and Analysis Pipeline

*Step 1:* Recording of Dyadic Interactions and Temporal Alignment. The first step is to record two participant’s eye gaze during social interaction. In our work, gaze is tracked using two Pupil Labs Invisible devices, which are mobile eye tracking glasses with one outward-facing camera to record the field of view, and infrared cameras in the frames to record the eye movement. Audio is also captured and stored together with video and gaze data on a cellphone. The Pupil Labs Invisible look and wear like classic wayfarer style glasses, which greatly increases user comfort and ecological validity.

A critical step for assessing dyadic gaze is that the data from both eye-trackers be aligned in time. We accomplish this via a sound marker (clap) that has a sudden, readily detectable onset, and gets recorded by both eye-trackers. Using the Pupil Labs exporter software, we identify the clap in the recordings and export all data subsequent to this common onset point.

*Step 2:* Gaze Location Identification and Target Identification. Once data was collected and synchronized, the raw data pulled from the phones was exported through Pupil Labs proprietary exporting software Pupil Labs Player, which provided gaze location via ‘X,Y’ coordinates over time. Now that we know where a participant is gazing, targets of interest in their field of view need to be identified, which in this case is the face of their conversational partner. We used the DensePose-Pupillabs module (https://github.com/pupil-labs/densepose-module) to determine the ‘X-Y’ coordinates of the face of the conversational partner for every frame of the outward facing camera. DensePose (**?**) can estimate the accurate head/face shape and the customized module combines the gaze and target location data for every frame and determine moments when participants are gazing at the face of the partner during the conversation, which gets stored as a CSV-file. We use this CSV as input to a custom-python program that restricts the data to focus on the face-gaze data and generate an equidistantly resampled time-line of annotations that indicate whether the face of the partner was looked at or not (coded as 1 and 0). Because our goal is to study social gaze, this pipeline is applied to data from each conversation partner.

*Step 3:* Merge Gaze Annotations from Interactants to Create Dyadic Social Gaze Tables. To combine the annotated data from each interactant, another python script reads in the individual gaze-annotated files and merges them based on the common timeline information. This yields a combined, tab-delimited file that includes timings, whether participant 1 looked at the face of participant 2 (subject1-looksAt-subject2), or vice versa (column: subject2-looksAt-subject1). This merged dyadic gaze-annotation file forms the basis for subsequent analysis of joint gaze.

(*Optional Step 4*: Add in Information about Speech Turns). To enable multimodal analyses of social communication, it is desirable to add in information beyond gaze alone, such as who is speaking and potentially even what is being said. Therefore, we expanded the core gaze-pipeline above to also include computerized annotations of speaker and listener behaviors. The sound recordings are diarized, providing information about who is talking (speaker01, speaker02) and the on/offset of talk-turns. This is accomplished via the PyAnnote (https://huggingface.co/pyannote/speaker-diarization) package. We then use a script to generate a tab-delimited file that matches the gaze-annotation timeline, holding on/off (1s and 0s) information about which person is speaking (columns: s1-speaking, s2-speaking). Finally, this file is merged with the gaze-annotations, yielding an output file that has 5 columns - time, s1-speaking, s1-lookingAt-s2, s2-lookingAt-s1, s2-speaking.

### Pilot Study: Capturing Social Gaze in Dyadic Settings

Before developing the actual gaze-analysis pipeline, we first conducted a pilot study. The purpose of this was to demonstrate that the mobile eye-trackers are precise enough to detect fixations at particular points in the visual field, especially the face, and capable of doing so at a normal conversational distance and with movement from both participants.

#### Approach

We will keep the description here rather short and provide further details in the Supplementary Materials. In brief, two participants were seated across from each other and each participant received different instructions via ear-phones to look at either colored marker points that were placed on the walls behind the opposing participant, or look at the face of the participant. This allowed us to test whether the eye-trackers can be used to detect attention (gaze) towards the instructed targets, whether we can detect gaze towards the interaction partner’s face, as well as instances of mutual gaze (when both partners were instructed to look at the other’s face).

#### Results

As shown in Figure 2 for an exemplary dyad, results confirmed that the mobile eye-trackers correctly tracked participants’ gaze towards the instructed targets under normal head movement conditions, including face-gazing and mutual face-gazing periods. These positive results were confirmed in a sample of 14 dyads (see Supplementary Materials). Furthermore, this study helped to optimize the eye tracking calibration and analysis routines. In sum, this provides proof-of-concept for using mobile eye-trackers to measure social gaze with sufficient precision.

**Figure 1:**
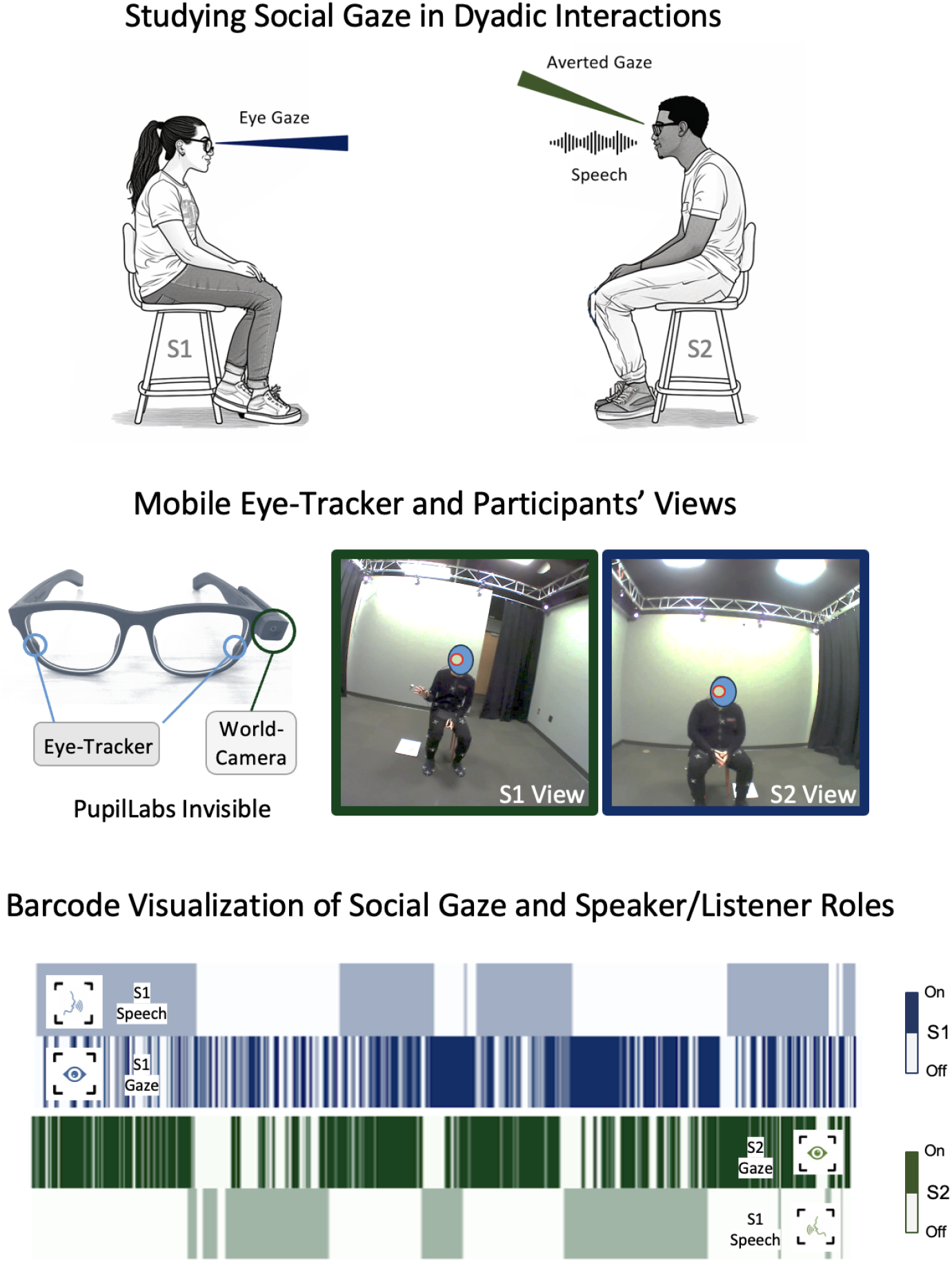
Study Results. A and B: Bar-Code plot illustrating gaze and speech events for both interactants for get-to-know and managerial interactions. C and D. Additional bar-code plots to illustrate both between-dyad variability as well as consistent patterns: Note, for instance the longer, more block-like duration pattern of speech turns in the managerial interactions. E and F. Results for all dyads. The mean for gaze represents the fraction of time participants spent gazing at their partner’s eyes/face over the total gaze time. The mean for speaking is the fraction of time talking over the total interaction time. A value of 0.5 would indicate they spoke or gazed exactly half of the interaction. There is a significant difference in time spent talking between the managers (sub001) and the employees (sub002) in the managerial role play interaction.

**Figure 2:**
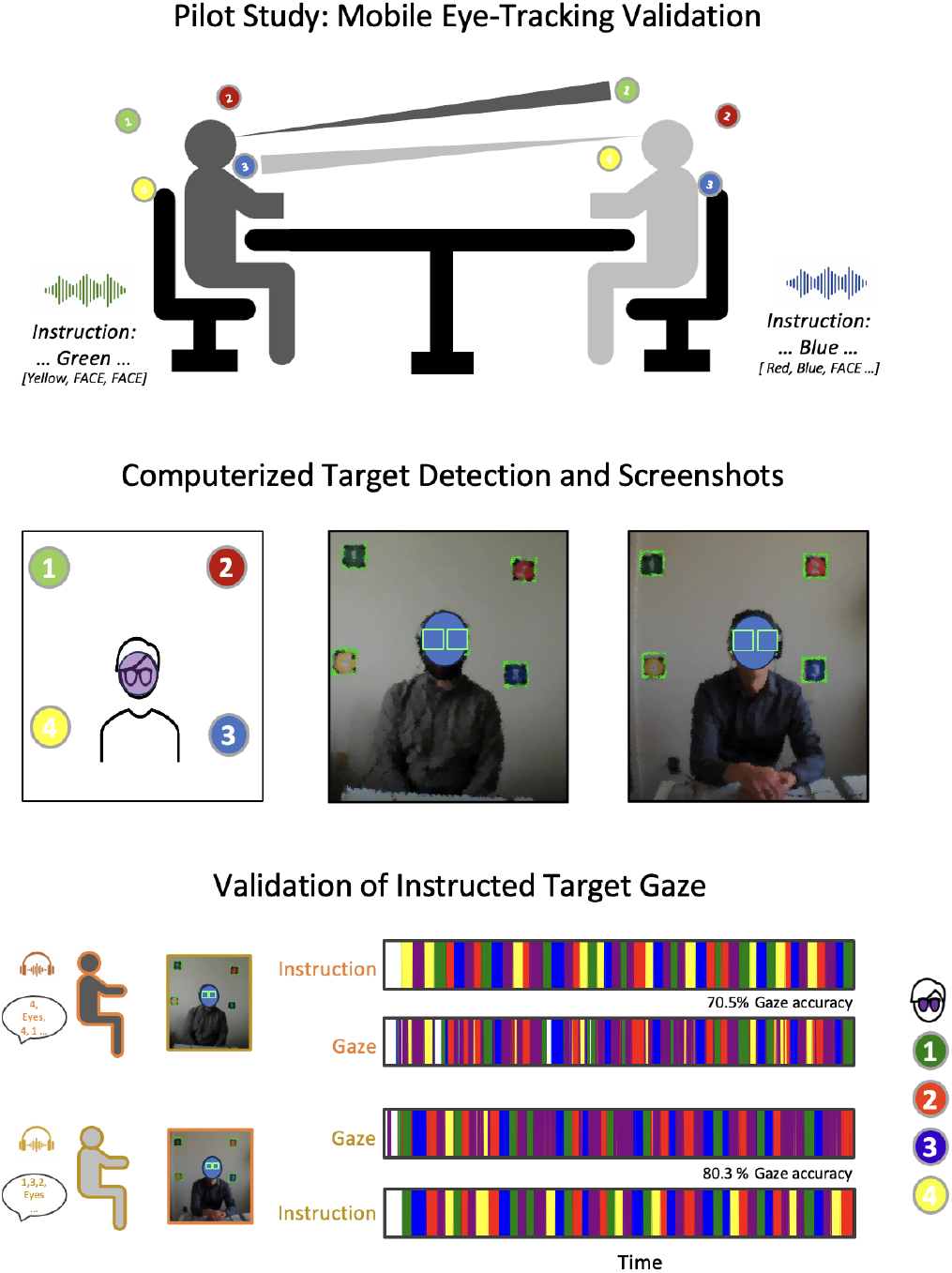
Pilot Study and Results for Validating the Accuracy of Mobile Eye-Trackers in a Naturalistic Dyadic Conversation Setting. A) Schematic of the setup: People are sitting opposed to each other and are instructed via earphones to look at particular positions, including the partner’s face (5). The bottom row shows results from one dyad demonstrating that gaze is tracked. (Note that differences between instructed and actual gaze locations are due to failure to immediately follow instructions rather than measurement imprecision). Images shared with permission.

### Validation Study: Automated Pipeline Assessment

#### Accuracy of a Computerized Social-Gaze-Analysis-Pipeline

Having established that the mobile eye-trackers can be used to detect instructed gaze targets in a conversational setting, we set out to validate the pipeline for face gaze detection in naturally running interactions.

#### Approach

Two expert coders used the electronic linguistic annotation tool (ELAN, Wittenburg et al. (2006)) to manually code gaze and speech data from three dyads who engaged in two short conversations each. We applied the new computerized pipeline to the same data and compared computerized results against ground truth as well as against the human annotations. Additionally, we also compared the main computerized analysis pipeline against a second, also computerized pipeline that relied on independent software algorithms (one using Python, the other Matlab; one using Pose/head detection, the other facial landmark detection). Methodological details and results can be found in the online Supplementary Methods.

#### Results

Comparing Computerized Gaze Analysis with Ground Truth. We find that computerized coding is highly accurate - reliable, objective and valid (see Figure 3 for exemplary results and Supplementary Materials for additional data and images of all coded interactions). When comparing the computerized (and human coding) with “ground truth” (original video with gaze coordinates superimposed), we find that the computerized method accurately detects all instances of face-gazing, and does so with extreme precision.

**Figure 3:**
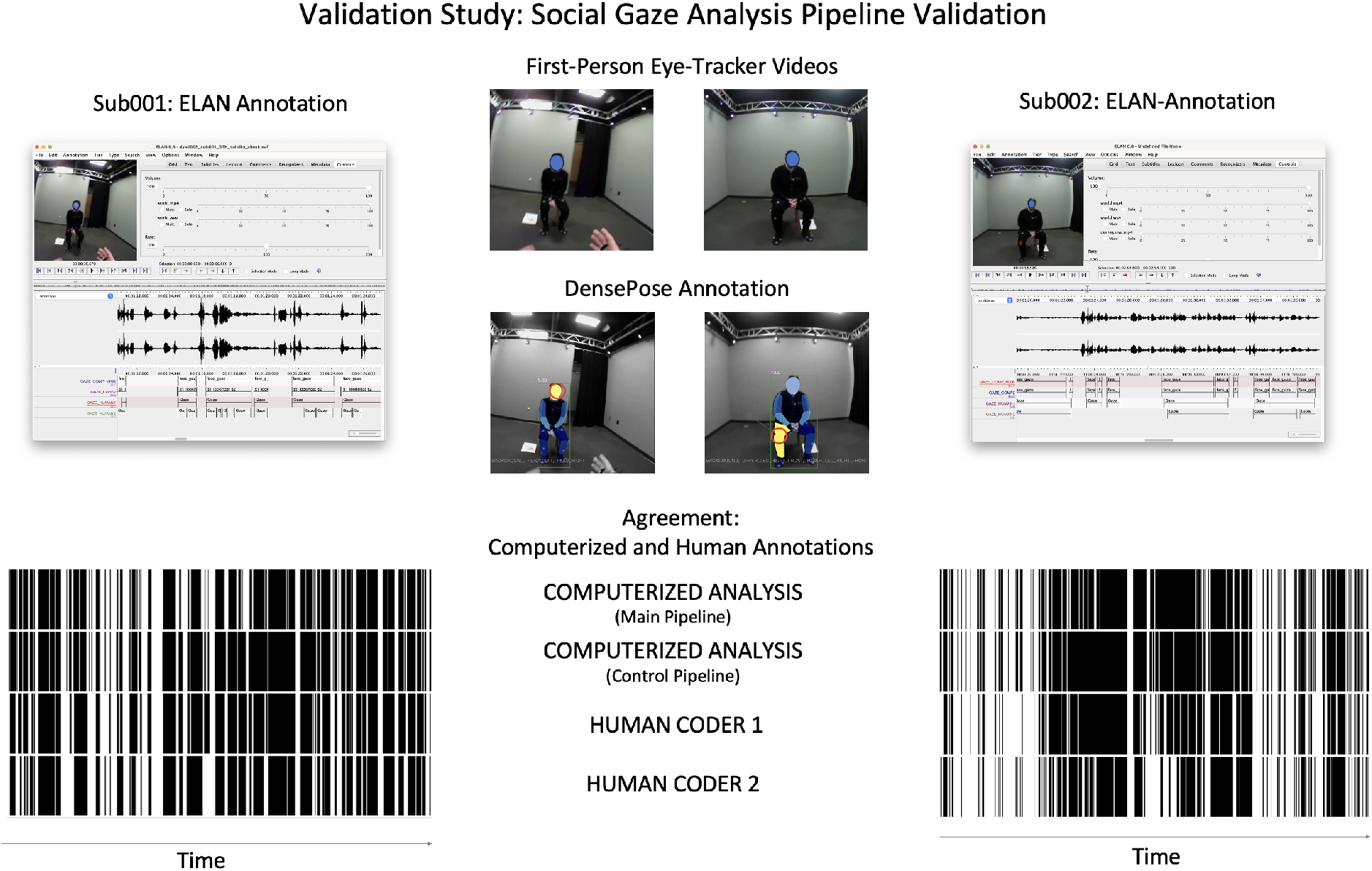
Validation Study Demonstrating the Accuracy of the Automated Social Gaze Analysis Pipeline. Top center screenshots illustrate the original recordings from the eye tracking cameras from two interacting participants. The corresponding screenshot from the DensePose package shows that one subject is gazing at the face of their partner, the other participant is currently looking at the partner’s knee. Left and right panels show screenshots of the ELAN tool, which was used to manually annotate the social gaze and import computerized annotations to compare them with ground truth. Bottom bar-codes demonstrate the agreement between the computerized pipelines and human coders. Visual inspection of the raw data revealed that human coders diverged particularly for short, transient gazes and differed slightly in timing (onset/offset). Images shared with permission.

Agreement among Human and Computer Coders. The two computerized face-gaze-identification methods show almost perfect agreement with one another (*κ* ¿ 0.95), with minimal deviations being attributable to the fact that the control pipeline only used a square bounding box around the face, whereas the main pipeline used the individually recognized face-area as the area of interest.

Agreement among the human coders is also good, but not perfect. On average, we find agreement between the two human coders to be *κ* = .58, sd = .16, although it could be as low as *κ* = .19 for an individual interaction. Human coders’ performance also showed good agreement betwith the computerized results *κ* = .64 (sd = .15), even slightly higher than between the two coders (because both are affected by error). See Supplementary Materials for detailed analyses of every interaction.

For the speech identification and diarization, the results were also positive (see Supplementary Materials for further details and result figures) and, in fact, even somewhat higher: Agreement among human coders amounted to *κ* = .9 on average with very high consistency across the twelve coded interactions (*sd* = .06, *min* = .77). Importantly, the agreement among human and computerized coding was high, amounting to *κ* = .81 (*sd* = .09; *min* = .68). Close inspection of the raw recordings and waveforms (ground truth) revealed no clearly discernible advantage for either computerized or manual coding. Differences between human coders and between human coders and the automated pipeline were largely attributable to the coding/non-coding of pauses/speech interruptions, which are a well-known source of ambiguity in speech analysis (Duncan and Fiske, 1977).

Taken together, these efforts validate the use of the computerized method for social gaze (and speech) analysis. The computerized method is more accurate and faster for gaze; for speech, both approaches exhibit similar performance, but again the computerized method excels at speech and scalability.

#### Computerized and Human Analysis of Social Gaze: Lessons Learned

These two studies demonstrate first the precision of mobile dyadic eye tracking and second the validity of computerized social gaze analysis. The advantages of the computer-based approach are clear and convincing - and they are the same as in other areas where computerized analysis has been profitably applied (Van Atteveldt et al., 2021; Paxton and Dale, 2013; Pouw et al., 2020; Araujo et al., 2020; Baldwin and Schmälzle, 2022): The computer precisely codes each and every frame, it does without any misses, temporal imprecision, or tiring out. Thus, the computerized pipeline can sift through large amounts of data at little cost. These measurement qualities strongly favor the computerized measurement of social gaze.

However, we note that close human inspection is still warranted - for two reasons: First, humans need to supervise the computer in order to detect drops in validity, or potential bias. For instance, our analyses show that the computers were able to detect faces of diverse participants in our college sample, but we did not yet check whether this would also apply to e.g. parent-infant dyads. Second, we still advise humans to code several dyads to familiarize themselves with the phenomenology of eye-contact and to potentially discover new patterns that have not yet been described in the literature. However, it is clear that coding hours of data on a frame by frame basis (and for each interactant, for several tasks, and for gaze and speech streams) is a daunting task that becomes quickly impracticable to perform at scale (Krippendorff, 2004; Riff et al., 2014).

### Gaze During Social Conversations in Two Settings

In this section, we apply the validated pipeline to an example study. In doing so, we aim to demonstrate the advantages of the computerized approach compared to previous research that relied on human observers and often only video recording (Thayer, 1969; Argyle et al., 1974; Levine and Sutton-Smith, 1973; Rutter et al., 1978; Kleinke, 1986), and embark on an exploration of gaze patterns and the social context variables that influence them.

Building on theoretical foundations and empirical findings laid by classic (Duncan and Fiske, 1977) and more recent (Ho et al., 2015), we characterize individual and dyadic gaze dynamics descriptively in terms of frequency, duration, and variability. Furthermore, we study the influence of social context variables (Foddy, 1978; Macdonald and Tatler, 2018) and the functional role of mutual gaze in turn-taking (Rutter et al., 1978; Wohltjen and Wheatley, 2021).

In brief, the data for this study comprise 78 interactions captured in 39 dyads who interacted over two tasks (citation: Database or NSF project). The primary manipulation is the social context of the interaction, which consisted of a relaxed get-to-know-you introductory conversation and a manager-employee interaction that established a power conflict. Previous gaze studies have shown that people exerting dominance have increased eye gaze when speaking (Thayer, 1969; Argyle et al., 1974; Exline et al., 1975; Kleinke, 1986). This is contrary to normal interactions where eye gaze is high during listening and gaze avoidance occurs while speaking (Argyle and Cook, 1976; Patterson, 1982; Kleinke, 1986; Wohltjen and Wheatley, 2021).

Another goal was to explore the relationship between eye gaze and the exchange of talk turns. In particular, previous work suggests that listeners tend to look more at speakers, with eye gaze serving as a backchannel activity to ensure the speaker of the listeners’ attention. However, the evidence is somewhat conflicting about whether and how eye gaze is used to initiate and end talk turns (Kendon, 1967; Levine and Sutton-Smith, 1973; Rutter et al., 1978; Wohltjen and Wheatley, 2021).

Thus, to measure and examine social gaze in this dataset, we applied the validated social gaze analysis pipeline described above. In the remainder, we briefly describe the study methods and then report the results of this example study.

#### Method

Participants. Participants for this study were undergraduates from a large Midwestern university who participated for course credits and/or monetary compensation in a larger study about dyadic social interactions. The sample consists of 39 dyads with complete data for both interactions, i.e. both tasks were completed and with eye-tracking data being available for all interactants and tasks (78 participants, or 156 individual interaction recordings). All participants were strangers to each other, provided written informed consent to the IRB-approved study.

Procedures. Every dyad first completed a get-to-know interaction, followed by a managerial interaction. The get-to-know conversation involved the two participants introducing themselves to each other and building rapport as they disclosed information. The scenario for the managerial task introduced a conflict between the manager and the employee, with the manager being in the higher social position, but both being motivated to achieve their specific goals, yet also motivated to achieve a resolution of the conflict. Both interactions took about five minutes.

Data Analysis. Data were analyzed using the computerized pipeline described above. Thus, we first aligned the individual participants’ recordings to the common sound onset, and then carried out the computerized routines to determine each participant’s gaze and speech behavior, yielding for every interaction a datasheet with four columns for the occurrence of face gaze and speech (i.e. whether sub001 or sub002 were speaking, and whether they were looking into each others’ faces, respectively). These datasheets formed the basis for subsequent analyses of eye-contact descriptives as well as inferential tests. Following the the seminal work of **?**, we developed code to automatically extract a list of 23 features that characterize either individual speech or gaze behaviors (e.g. total-time-speaking, fraction-time-speaking, total-time-gazing-at-partner, fraction-time-gazing etc.) as well as dyadic metrics (e.g. fraction-of-conversation-spent-making-eye-contact; average-duration-eye-contact; average-eye-contacts-per-minute). We then characterized all 78 interactions on all of those measures and compared them across the two tasks (39 get-to-know vs 39 managerial).

## Results and Discussion

The descriptive statistics shown in Table 1 and the bar-code plots in Figure 4 provide insights into the amount of time participants spent talking, eye gazing, and how conversational context affected them. Perhaps the most salient visual observation is that the average length of talk-turns appeared to increase from the get-to-know to the managerial task: Indeed, in the get-to-know interaction, many dyads spend considerable time with typical back-and-forth chit-chat, whereas the managerial task typically started with the boss (sub001) explaining the reasons for calling the employee to the meeting, followed by longer turns in which both parties tried to advance their goals.

**Table 1:** Means and Standard Deviations for Individual and Dyadic Gaze and Speech Features across social tasks. Note that for the managerial task, S1 had the role of the boss, and S2 the employee.

|  | Get-To-Know |  | Managerial |  |
| --- | --- | --- | --- | --- |
|  | mean | sd | mean | sd |
| total_number_S1_speaking | 28.08 | 11.21 | 14.21 | 10.55 |
| total_number_S2_speaking | 26.38 | 10.27 | 14.36 | 11.12 |
| total_number_S1_lookingAt_S2 | 92.31 | 62.35 | 85.05 | 60.4 |
| total_number_S2_lookingAt_S1 | 82.36 | 45.54 | 77.74 | 47.64 |
| average_length_S1_speaking | 5.45 | 2.86 | 10.98 | 9.59 |
| average_length_S2_speaking | 4.88 | 2 | 6.65 | 3.92 |
| average_length_S1_lookingAt_S2 | 2.59 | 1.96 | 2.42 | 1.66 |
| average_length_S2_lookingAt_S1 | 2.72 | 2.53 | 2.5 | 2.16 |
| fraction_time_spent_S1_speaking | 0.58 | 0.12 | 0.54 | 0.11 |
| fraction_time_spent_S2_speaking | 0.52 | 0.12 | 0.37 | 0.1 |
| fraction_time_spent_S1_lookingAt_S2 | 0.7 | 0.19 | 0.71 | 0.18 |
| fraction_time_spent_S2_lookingAt_S1 | 0.71 | 0.25 | 0.68 | 0.26 |
| S1_GazingFraction_When_Speaking | 0.63 | 0.2 | 0.65 | 0.21 |
| S2_GazingFraction_When_Speaking | 0.64 | 0.25 | 0.61 | 0.25 |
| S1_GazingFraction_When_Listening | 0.81 | 0.2 | 0.82 | 0.2 |
| S2_GazingFraction_When_Listening | 0.77 | 0.27 | 0.75 | 0.28 |
| dyadic_eye_contact_count | 129.13 | 60.66 | 121.33 | 71.32 |
| daydic_average_eye_contact_duration | 0.89 | 0.48 | 0.85 | 0.46 |
| daydic_number_eye_contact_per_minute | 33 | 11.01 | 34.4 | 13.07 |
| daydic_percent_eye_contact | 0.48 | 0.21 | 0.47 | 0.21 |
| daydic_fraction_speaking_silence | 0.04 | 0.03 | 0.11 | 0.07 |
| daydic_fraction_covered_speaking | 0.96 | 0.03 | 0.89 | 0.07 |
| daydic_fraction_overtalk | 0.13 | 0.06 | 0.02 | 0.02 |

**Figure 4:**
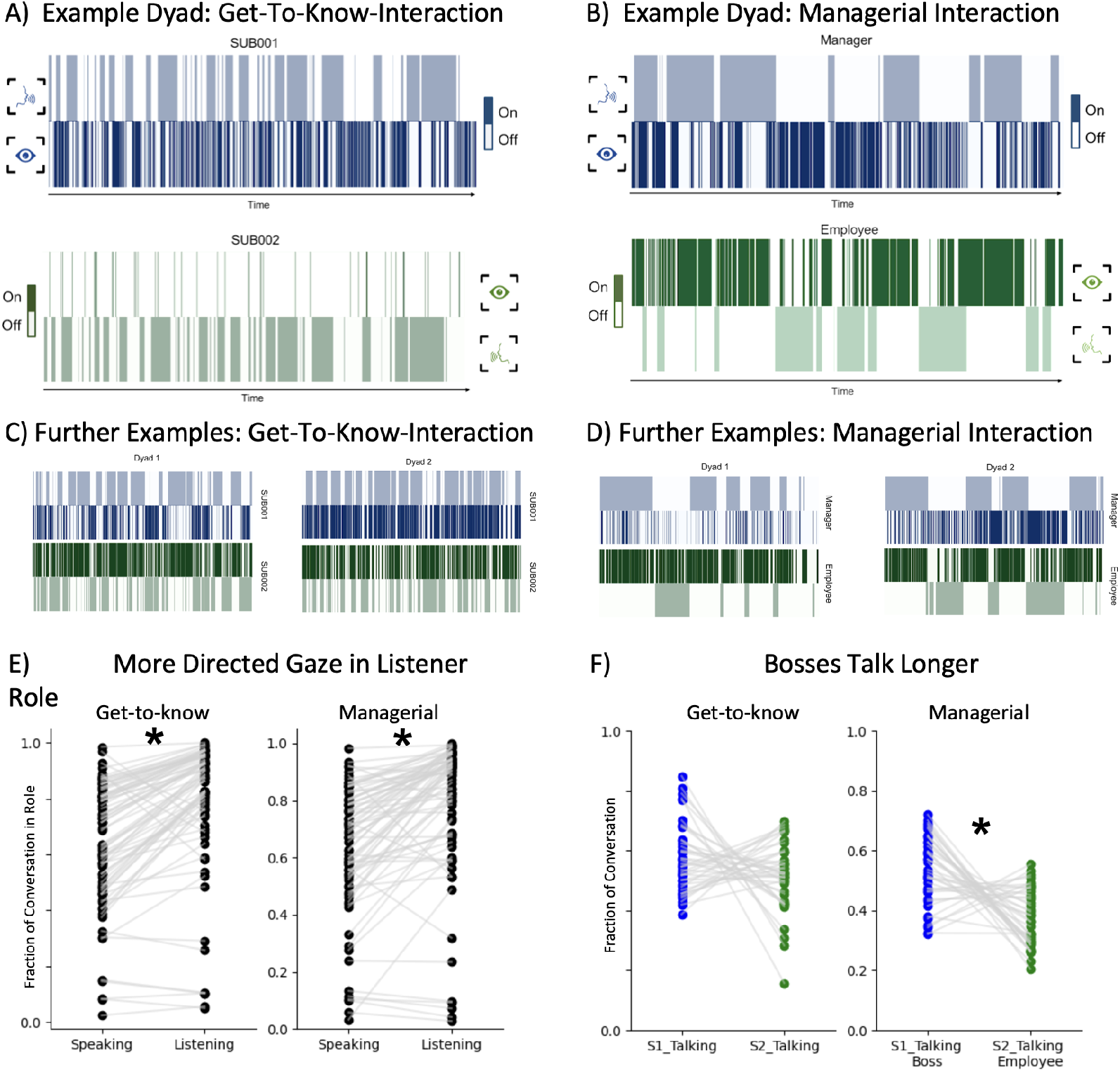
Study Results. A and B: Bar-Code plot illustrating gaze and speech events for both interactants for get-to-know and managerial interactions. C and D. Additional bar-code plots to illustrate both between-dyad variability as well as consistent patterns: Note, for instance the longer, more block-like duration pattern of speech turns in the managerial interactions. E and F. Results for all dyads. The mean for gaze represents the fraction of time participants spent gazing at their partner’s eyes/face over the total gaze time. The mean for speaking is the fraction of time talking over the total interaction time. A value of 0.5 would indicate they spoke or gazed exactly half of the interaction. There is a significant difference in time spent talking between the managers (sub001) and the employees (sub002) in the managerial role play interaction.

Zooming in specifically on the individual gaze and dyadic eye-contact metrics revealed interesting results. First, we find that the average duration of directed social gaze (defined as periods in which one partner is gazing at the other partner’s face, irrespective of whether this is reciprocated) is around ca. 2.3-2.7 seconds, which aligns with prior reports in the literature. Also, it can be seen that the fraction of gazing is higher when speaking compared to listening (e.g. 0.63→0.81; 0.65→0.82; 0.64→0.77; 0.61→0.75 for speakers 1 and 2 in the get-to-know and managerial tasks, respectively).

Statistical comparisons confirmed that the fraction of time spent gazing at the interaction partner differed depending on the speaker/listener role. Specifically, in the get-to-know task the fraction of gaze while listening (Listener-Eye-Gaze, LEG) was *mean*_*LEG*_ = .79 (*sd* = .24) compared to *mean*_*SEG*_ = .63 (Speaker-Eye-Gaze, *sd* = .22), which is a significant difference (*t* = 9.5; *p* < .001). Almost identical values were observed for the managerial interaction (*mean*_*LEG*_ = .78; *sd* = .25; *mean*_*SEG*_ = .63, *sd* = .23, *t* = 8.02; *p* < .001), providing an internal replication.

Periods of eye-contact are comparatively shorter - slightly less than a second on average, which again matches prior observations. We find that on average, about 50-percent of the time during conversation are spent making eye-contact. This aligns with the conversational nature of the tasks, which also have about equal fractions of speaking. Note, however, that periods of directed social gaze (i.e. one partner looking at the other) and periods of directed speaking do not map onto each other in a one-to-one fashion (see Figure 4), but rather gaze behavior is interdigitated with speaking/listening roles, underscoring its functional role for interaction regulation (e.g. Patterson, 1981). We did not observe significant differences in the time spent making eye contact between tasks.

Importantly, it should not go unmentioned that these averages and group statistics, which are in line with prior reports, provide aggregate summaries of phenomena that exhibit marked individual differences. For instance, as can be seen in Figure 4 (bottom panel), some dyads exhibited marked imbalances in both eye-contact as well as speaking fractions. To delve into this, we additionally computed not only standard deviations for all of these scores, but also the coefficients of variation (mean/sd) for all metrics. Inspecting those metrics revealed marked between-dyad differences, for example, for the speed and frequency of turns as well as several eye-contact metrics.

Statistical comparisons also confirmed that the fraction of time during which the manager talked (compared to the employee) increased markedly: Comparing the speech-fraction between the manager vs. employee revealed a significantly higher rate for the manager (*mean*_*speech-fraction-manager*_ = .54, *sd* = .11) and the employee (*mean*_*speech-fraction-employee*_ = .37, *sd* = .09; *t(38)* = 5.12, *p* < .001). Furthermore, there was also a difference between the employee’s speech share during the managerial task when compared to the same person’s speech behavior during the preceding get-to-know task (i.e. serving as an intraindividual baseline), *t(38)* = 6.23, *p* < .001. Several other conversational metrics confirmed these differences between tasks. For instance, the fraction of the conversation spent in silence was much higher in the managerial task (*t(38)* = 6.52; *p* < .001; and - correspondingly - the occurrences and rate of periods during which the interactants both spoke or one of them spoke differed). As can be seen from the figures, the instances and durations of speech turns also varied, i.e. fewer but longer speech turns by both speakers as well as employees (number, S1: *t(38)* = 6.71, *p* < .001; S2: *t(38)* = 6.12, *p* < .001; average length: S1: *t(38)* = -3.94, *p* < .001; S2: *t(38)* = -2.41; *p* < .05).

To appreciate these results, it is helpful to interpret them in light of classical research. Almost 50 years ago, Duncan and Fiske (1977) published a seminal book titled ‘Face-to-Face Interaction. Research, Methods, and Theory’. This gem of non-verbal communication scholarship reports a careful and exhaustive study of dyadic conversations conducted under “as close as possible to natural conditions”. The authors coded over 30000 acts from about 40 dyads during short videotaped conversations. The resulting 380 pages with dozens of tables describe a monumental project, which spanned several years and involved many coders. The current study, though less ambitious in its goal (Duncan and Fiske’s ambition was to develop an all-encompassing theory of conversational behaviors, which is presented in the later part of the book), features a comparable amount of data (ca. 40 dyads) recorded and analyzed with updated methods. Although we focus only on gaze and speech signals, this serves to showcase the enormous potential of the current approach.

In fact, as the results in Table 1 demonstrate, we detect several classical findings in gaze and turn-taking research. For instance, directed gaze (at the face) takes about 3 seconds, and this metrics was discovered similarly in both subjects as well as across both tasks. That durations of eye-contact are shorter than individual gaze-turns is not only mathematically necessary, but also aligns with the fact that too-long eye-contact can become uncomfortable, especially in conversations between unacquainted strangers as the current one. Overall, the promise of this approach lies in its rigorously quantitative nature, which now, given that the measurement and analysis challenges are solved, can usher in a new era of social signal processing (Burgoon et al., 2017; Bredin et al., 2020). Of course, the current analyses and metrics have barely scratched the surface of the richness of these data. For instance, it will be worthwhile to examine questions like the dynamics of eye-contact markers over time (how do they evolve or fluctuate over a conversation or developing relationship?), their intrapersonal stability (e.g. is the social gaze behavior like a fingerprint that could e.g. be detected when exhibited by a person’s avatar in VR?) as well as interindividual and cultural differences. This will allow researchers to mine the rich patterns of social gaze, promising fruitful insights into human interaction, social cognition, and the biological underpinnings and social consequences of social gaze (Hessels et al., 2023; Tomasello et al., 2005).

## Main Discussion

Social gaze, and in particular the ensuing periods of eye contact, is a critical component of human communication. However, its intricacies have long remained somewhat enigmatic and difficult to study. The current work harnesses innovative mobile eye tracking technology to delve into the minutiae of how social signals are exchanged during dyadic interactions. We introduced, validated, and showcased methods that can propel the study of eye gaze in communication to a new level. Below, we first highlight the methodological contribution and then discuss insights gained from the example study. We then cover the theoretical and applied implications, remaining limitations, and avenues for future research.

### Discussion of Main Results

First, the methodological contribution of this work is that it provides strong support for the use of mobile eye-trackers and computerized methods to assess social gaze in interactions. As we have shown above, the mobile eye trackers are sufficiently precise and the automated pipeline is objective, reliable, and valid. Together with the well-known benefits of automated coding - scalability and cost-effectiveness - this recommends wider adoption in the field of communication.

In addition to validating this pipeline, applying it to an example dataset offered fine-grained insights into nonverbal social interaction patterns. We replicated some well-known statistics that characterize individual and dyadic social behaviors. Furthermore, by making our code for gaze analysis and gaze-statistic quantification publicly available, other researchers can build on and expand on this work. Now that the validity of the computerized analysis has been demonstrated, the focus can shift on the task of acquiring as-clean-as-possible data and from a wide variety of communication situations. The former (acquiring clean data) may perhaps seem obvious, but is far from trivial given the complexity of dyadic research and the measurement equipment (Huskey, 2024). For instance, if only one of the dyadic eye-trackers fails, the entire dyad has to be discarded; similarly, if only one microphone collecting speech data is covered or records suboptimal data, this affects the speech diarization; or, if the gaze calibration is off, accuracy of any subsequent analysis (computerized or human) will suffer.

In sum, while the current data strongly supports the use of the pipeline presented here, the data acquisition could likely be improved. We also envision that future mobile eye-trackers will be more error-tolerant and feature added capabilities, such as automatic calibrations/re-calibrations. Finally, as noted above, we still favor a human-in-the-loop approach for quality assurance as well as to equip new scholars of nonverbal communication with the necessary expertise of social gaze behaviors.

### Avenues for New Research - Theoretical and Applied

Theoretical studies of Social Interaction Phenomena. With mobile eye-trackers enabling naturalistic social interaction research, and flexible analysis pipelines like the one suggested here, we are about to enter a new era of nonverbal communication research. This enables us to study a plethora of aspects relating to gaze - revisiting, replicating, and expanding on the groundwork laid by classical research (Burgoon and Le Poire, 1999; Argyle and Cook, 1976; Duncan and Fiske, 1977). Questions that can bow be tackled range from simple ones like the impact of physical characteristics (e.g. attractiveness) to a variety of subfields within interpersonal communication. For instance, we see much potential for this work in the area of family communication, interpersonal conflict, social influence, as well as in health-related settings (example citations).

While we deliberately kept this study within a laboratory setting to control situational factors, it is very feasible to further release the grip of experimental control and apply this work to more naturalistic settings, such as walking-and-talking conversations (where visual attention is on the street to avoid falling, but eye-contact matters), business meetings, or public speeches given at social gatherings and academic conferences (LeFebvre et al., 2021).

Another area for expansion lies in the range of social signals captured. As said, we focused here mainly on eye-contact and the closely related speech-turns, but comparable work exists on e.g. facial expressions, body language, paraverbals, and gestures (Burgoon et al., 2017). Thus, the multimodal integration of all of these channels is now within reach.

Potential for Applied Impact. One intriguing area for practical studies of eye gaze lies in how the social attention it signals impacts conversation. For instance, it is nowadays easy to find oneself trying to have a serious conversation with someone who is looking at their phone. This so-called “phubbing” (phone snubbing), which can lead to negative attributions and relationship satisfaction (Beukeboom and Pollmann, 2021), could now be studied empirically and integrated with social cognition research.

Relatedly, it has long been shown that positive and effective patient-physician communication improves health outcomes (Stewart, 1995). Research has shown that people avoid seeking medical treatment due to communication issues and concerns whether the physicians care about the patient (Taber et al., 2015). Characterizing and potentially intervening to improve gaze during patient-physician interactions provides relatively simple lever to improve communication, conveying a sense of caring and active listening on part of the physician, which inturn may lead to better health outcomes (Jongerius et al., 2020). Last, but certainly not least, we see large potential for applied impact in the context of virtual and computer-mediated interactions. Indeed, popular online-meeting platforms like Zoom suffer woefully from a lack of eye-contact. This has consequences for feelings of mutual connection, but also the ability to convey and contextualize nonverbal messages, or to use eye-gaze for turn-taking orchestration as studied above. Along similar lines, as we are about to enter the Metaverse, i.e. using virtual and extended reality for communication. In these contexts, it will be extremely important to more fully incorporate eye gaze, as otherwise the avatars or embodied agents will remain socially stale and deficient. In face, this has already led to leading VR manufacturers to include eye-trackers into their devices, and we can see a growing interest in research on avatars and embodied conversational agents to also focus on gaze.

With this in mind, the current work underscores the importance of the basic scientific sequence from description to explanation to prediction and control: Only if we are able to precisely characterize the phenomenon (e.g. eye contact, its duration, its variability, and so forth) will we be able to develop theories that explain its generative mechanisms. Then, if we are able to understand the governing principles of how eye-contact works, we will be able to predict whether it will occur, and ultimately may even intervene in the form of assistive technologies or socially adept AI-agents/robots (Georgescu et al., 2014; Roth et al., 2015).

### Limitations

While this study offers a methodological advancement, like all research, it has limitations. One limitation it shares with all bio-behavioral/apparative approaches is that equipment malfunctions are still a significant issue. In fact, for dyadic research this issue becomes all the more severe as the number of potential failures multiply and interact. Second, it could be seen as a limitation that one needs two devices, which are not all too expensive, but still expensive enough to require external funding. However, compared to e.g. the funds it nowadays takes to acquire high-quality online data from survey panes, this investment seems of a similar magnitude. Unfortunately, given the low precision, the work presented here cannot yet be used with e.g. phone-based or webcam-based trackers, but technological advances may eventually overcome this limitation. Another, somewhat more theoretical or definitional limitation refers to the distinction between narrow definitions of eye-gaze (pupil-to-pupil) vs. the wider definition of face gaze. In the current work, we focused on the latter, which is in line with other approaches and seems appropriate given the role of face gaze as an indicator of social attention, but still worth pointing out (Jongerius et al., 2020). Finally, although our approach is automatized, there is still some limited hand-intervention necessary to assure data quality, align dyadic recordings, and disambiguate speaker roles in the diarization process. We expect, however, that advances in AI will soon be able to also automate this piece, leaving researchers more room to focus on the theoretical mechanisms and functions of eye gaze.

## Conclusion

The moment when eyes meet is a pivotal moment for social communication. For instance, the first relationship building between parents and their kids occurs via gaze, but gaze also plays a fundamental role in all spoken conversations. Eye gaze has long been of interest to scholars, but has been plagued and stymied by the difficulty of efficiently measuring gaze, especially in naturally interacting dyads. This study achieved the goal of accurately assessing eye gaze and establishing an analysis pipeline for studying gaze in dyadic interactions. The ‘tip of the iceberg’ is an apt metaphor for where gaze research stands within the field of communication. This work represents a crucial step into peering under the water.

## Data Availability Statement

Data and code are available at osf.io/mvpf5

## Acknowledgements

We thank the participants of the study. We thank the developers of the densepose-pupillabs software package as well as the python packages used in this work.

## Funding Statement

Support for this research was provided by the National Science Foundation via the grant IIS-1907807

## Supplementary Materials

### A Pilot Study: Supplementary Methods and Results

The experimental paradigm was created that maximized ecological validity of a casual conversational environment while maintaining rigorous test of the eye tracking glasses. Two participants were seated across from each other in a natural conversational setting (See Figure 2 in main text), and each participant received different instructions to attend to certain targets in their field of view. The instructional targets were four stationary numbers on the wall, with each printed on uniquely colored backgrounds, which correspond to the colors in the barcodes in Supplementary Figure S1, and the fifth target was the face of their partner (which was dynamically moving). This instructional phase went on for two minutes with each participant receiving 50 targets, so target gaze time averaged two seconds. After the instruction phase ended, a two minute conversational phase was conducted. Results on this conversation will not be reported here, as the emphasis of the pilot study was to validate the hardware. In all, 17 dyads recorded, 3 discarded, leading to 14 dyads, 28 instructed recordings, being analyzed.

**Supplementary Figure 1:**
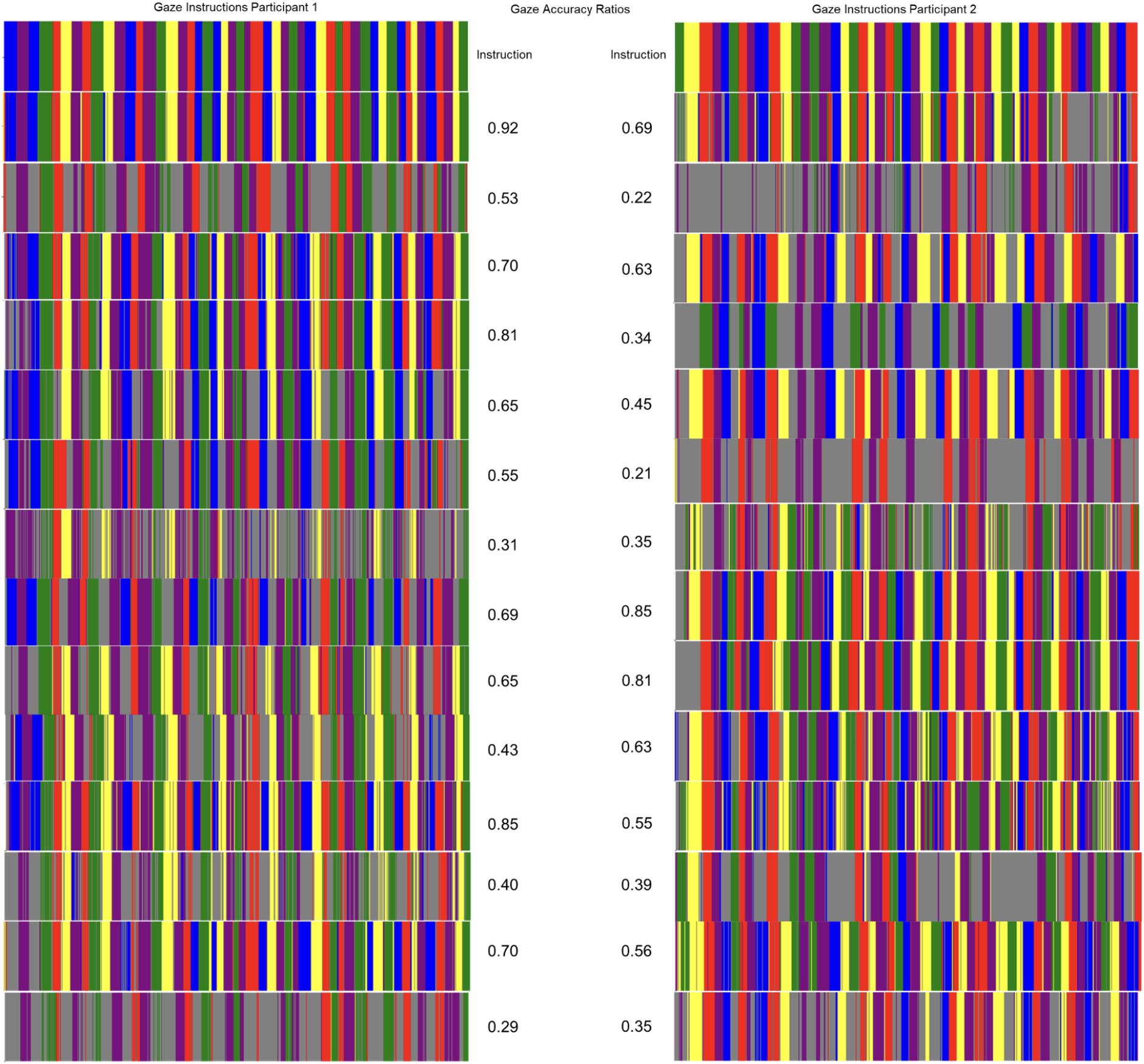
Pilot Study and Results for Validating the Accuracy of Mobile Eye-Trackers in a Naturalistic Dyadic Conversation Setting. Accuracy scores were given to each participant by creating a ratio gaze-on-target/total-gaze (mean accuracy 55.4-percent).

The analysis pipeline for this validation differed from the proposed Main Pipeline detailed in Supplementary B1 Methods. It utilized the custom MATLAB code that enabled us to track regions of interest (four stationary targets on the wall) and used the MTCNN-face-detection package (Pinkney, 2022) to track the faces of participants.

As seen in supplementary Figure S1, accuracy scores were created for every participant based on the time they gazed at the target corresponding to the instruction given. The average across all the participants was 55.4-percent. However, when diving into the video recordings, it was evident that participants struggled to identify the correct target, often bouncing between multiple targets before settling onto the correct one. Ultimately this result was promising that the eye trackers accurately and reliably tracked the gaze of the wearers.

### B Validation Study: Supplementary Methods and Results

Human Coding Procedures. Two expert coders manually coded all instances of eye gaze and speaker/listener roles in the interaction recordings. The data for this validation study came from three freely interacting dyads who had two short conversations, an initial get-to-know conversation followed by a managerial role play. Thus, with three dyads comprising two partners each, and two conversations from each dyad, this yielded twelve individual to-be-coded datasets.

The two expert coders used the ELAN tool (Wittenburg et al., 2006) to independently annotate face gaze or speaking roles, respectively. Specifically, each dataset’s eye-tracking video (containing the field of view with the overlaid gaze point) was loaded into ELAN and instances where the gaze point fell within the face were manually marked as accurately as possible.

Analogously, the sound-wave (also recorded from the eye-tracking device along with the video stream) was loaded, listened to, and moments in which the participant was speaking were annotated based on the sound’s waveform and by listening in on the sound to determine it came from the speaker (and not the other participant).

This coding process is very laborious, slow, and potentially error prone. Annotating a single few-minute conversation in this fashion took about 30-60 minutes. Thus, annotating the entire validation dataset (2 coders, each coding 3 dyads with 2 people, for 2 tasks, and annotating the 2 channels of gaze and speech) quickly became a painstaking task that took about a week and led to coder wearout, performance drops, and high personnel costs.

Computerized Coding Procedures. Main Pipeline (Densepose) For the computerized coding, we carried out the automatized pipeline described above for each dyad and conversation. We then imported the results to ELAN software for visual inspection along with the human coders’ annotation. Although setting up and validating this pipeline was carried out over a longer period, now that the tools exist and are vetted, it can be used to annotate entire datasets within less than a day assuming that data are in the appropriate format and named consistently (after the BIDS standard).

Control Pipeline (MTCNN). In addition to using the proposed computerized pipeline (Main Pipeline), we set up a second pipeline that also made use of computerized analysis, but relied on different, independent tools. Specifically, this secondary pipeline was based on MATLAB instead of Python code, and it leveraged the MTCNN-face-detection package (Pinkney, 2022) instead of the DensePose package. Thus, we essentially wanted to have an independent computerized software carry out the same task - akin to having two human coders.

As can be seen in Supplementary Figure S2, computerized and human-coding show good agreement. However, human coders sometimes exhibit stretches of inattention/perseverance, failing to code smaller gaze interruptions. Also, coders may differ in terms of the minutiae of onset times, especially given the fast frame rate of 30 fps.

**Supplementary Figure 2:**
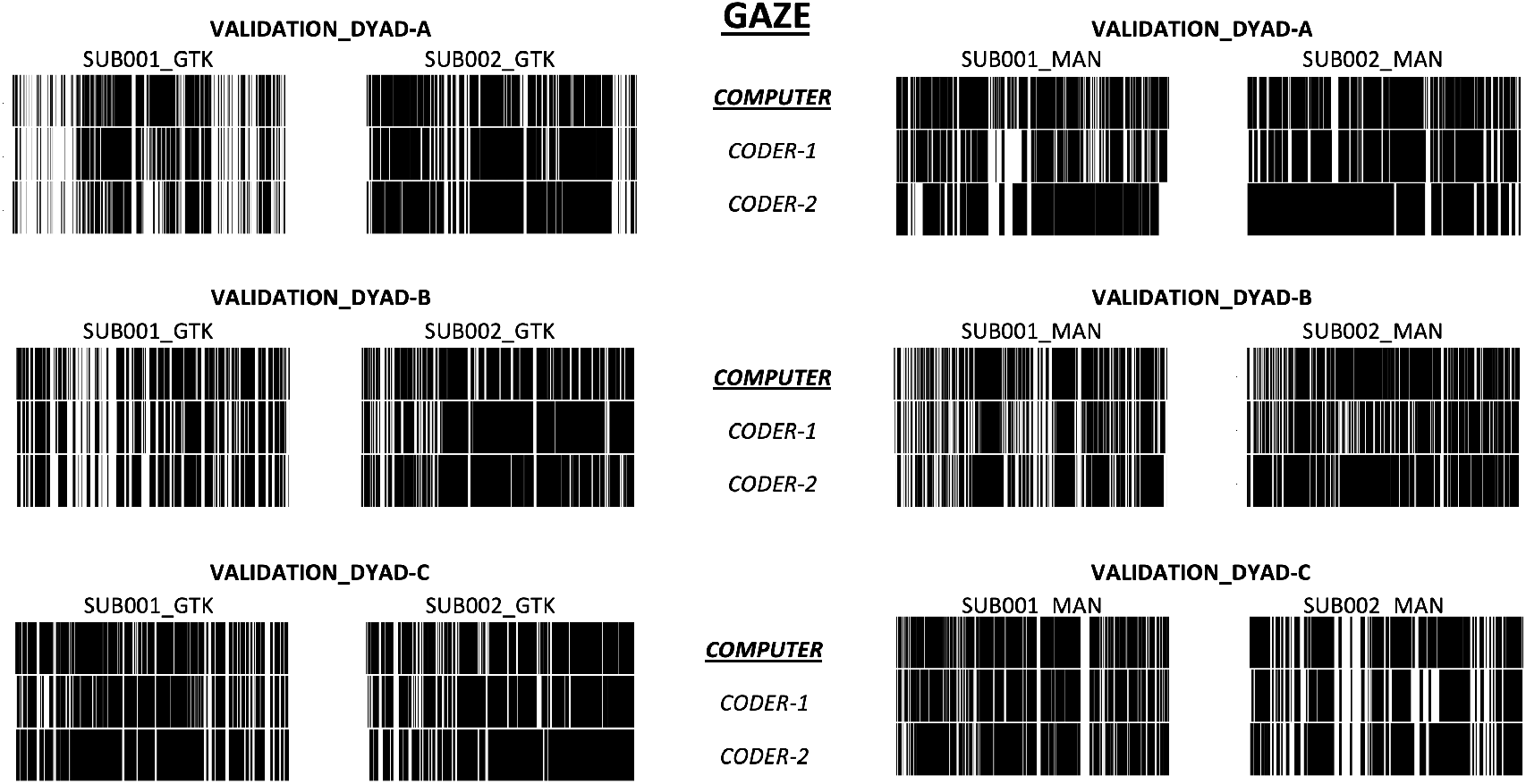
Gaze-Barcodes for the three validation study dyads for both tasks (Get-to-Know and Managerial Interaction

**Supplementary Figure 3:**
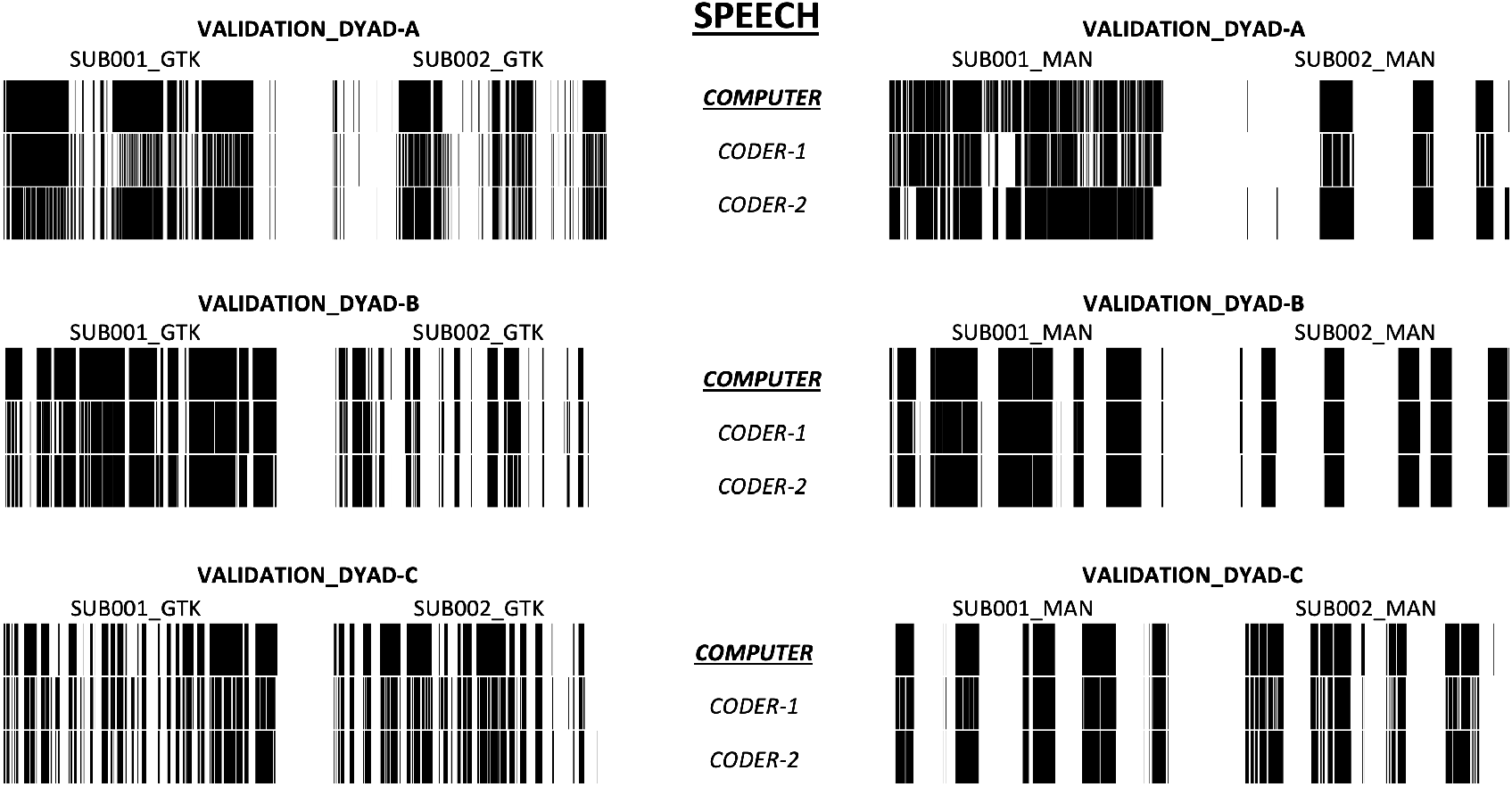
Speech-Barcodes for the three validation study dyads for both tasks (Get-to-Know and Managerial Interaction). Note the far more patterned, complementary structure (speaker-listener-roles) of these plots compared to the gaze-behavior shown in Supplementary Figure 2

**Supplementary Figure 4:**
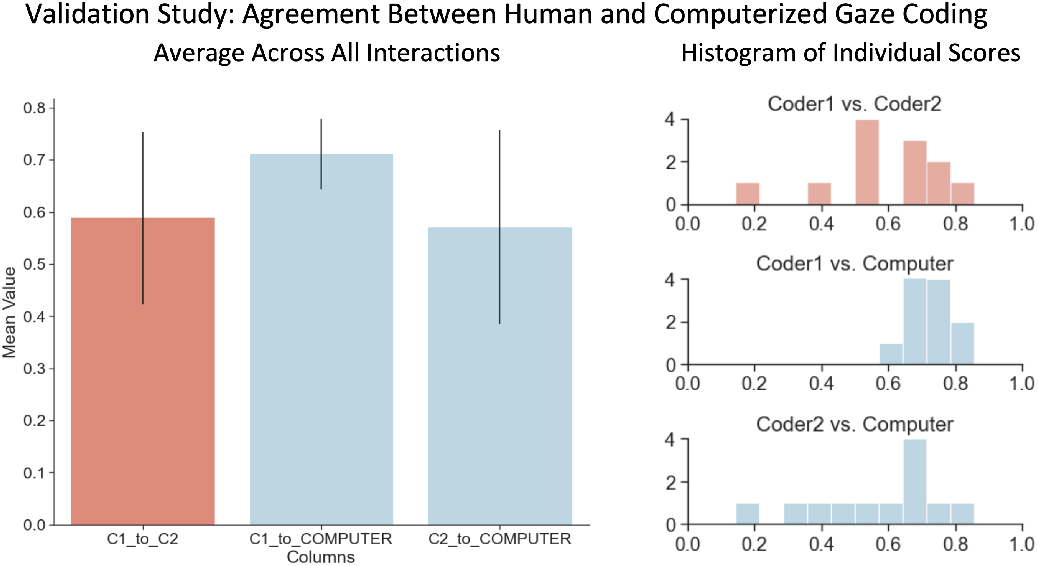
Mean agreement for gaze-coding: Human-to-human and human-to-computer-annotation comparison. Left: Mean and standard deviations across all 12 interactions. Right: Histogram of individual results for all 12 coded interactions.

**Supplementary Figure 5:**
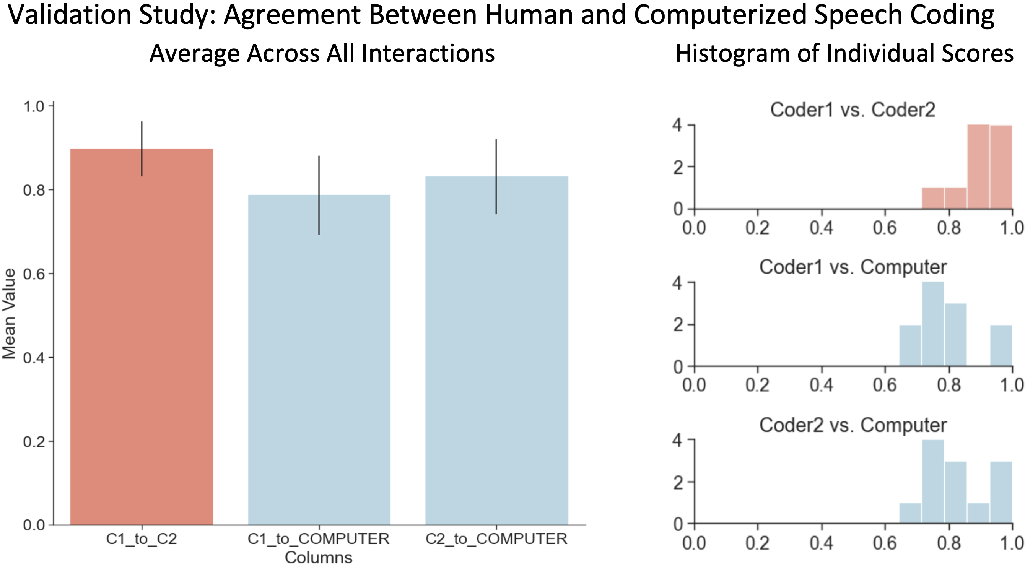
Mean agreement for coding of speech (speaker/listener) periods: Human-to-human and human-to-computer-annotation comparison. Left: Mean and standard deviations across all 12 interactions. Right: Histogram of all individual coding results.

## Notes

### Competing Interest Statement

The authors have declared no competing interest.

